# SGCRNA: Spectral Clustering-Guided Co-Expression Network Analysis Without Scale-Free Constraints for Multi-Omic Data

**DOI:** 10.1101/2025.04.27.650628

**Authors:** Tatsunori Osone, Tomoka Takao, Shigeo Otake, Takeshi Takarada

## Abstract

Weighted Gene Co-expression Network Analysis (WGCNA) is among the most widely employed methods in bioinformatics. WGCNA enables the identification of gene clusters (modules) exhibiting correlated expression patterns, the association of these modules with traits, and the exploration of candidate biomarker genes by focusing on hub genes within the modules. WGCNA has been successfully applied in diverse biological contexts. However, conventional algorithms manifest three principal limitations: the assumption of scale-free topology, the requirement for parameter tuning, and the neglect of regression line slopes. These limitations are addressed by SGCRNA.

SGCRNA provides Julia functions for the analysis of co-expression networks derived from various types of biological data, such as gene expression data. The Julia packages and their source code are freely available at https://github.com/C37H41N2O6/SGCRNA.

## Introduction

A gene co-expression network represents relationships among genes by depicting those with correlated expression levels as a network graph. In this approach, genes are treated as nodes, with edges connecting nodes that exhibit correlated expression levels. In Weighted Gene Co-expression Network Analysis (WGCNA), correlation coefficients are weighted to emphasise the strength of association[1]. Since its introduction, WGCNA has been employed in numerous biological studies. As of early April 2025, a search for “WGCNA” on PubMed yields more than 8,000 publications. For instance, WGCNA has been utilised in a study investigating the impact of microplastics on fish[2]. In this study, WGCNA was applied to RNA expression data from different exposure types to elucidate how ingestion through diet and respiratory filtration via the gills affect hepatic metabolism. In another investigation, WGCNA was employed to identify molecular markers[3]. This research applied machine learning models and WGCNA to peripheral blood proteomes to explore molecular markers for the rapid and accurate diagnosis of influenza virus infection. Beyond transcriptomic and proteomic analyses, WGCNA has also been applied to DNA methylation datasets. For example, WGCNA has been applied on DNA methylation patterns in the brains of individuals with Alzheimer’s disease[4].

In this manner, numerous studies have identified genes essential for biological mechanisms through WGCNA. Consequently, various tools have been developed to enhance the applicability and usability of WGCNA. For example, PyWGCNA, a Python-based implementation of WGCNA, has been developed[5]; CoExp has rendered WGCNA accessible via a web-based interface[6]; hdWGCNA has been optimised for single-cell and spatial transcriptomic analyses[7]; CWGCNA has facilitated causal inference within co-expression networks[8]; and TPSC has employed spectral clustering in place of hierarchical clustering[9].

Despite its extensive use, WGCNA possesses certain limitations. Firstly, it relies on the assumption of a scale-free topology[10]. Historically, many networks, including but not limited to co-expression networks, have been regarded as exhibiting scale-free properties. A scale-free network is characterised by a degree distribution that follows a power law, wherein a few nodes possess a large number of connections, while the majority have relatively few. However, recent studies suggest that scale-free networks are less prevalent than previously assumed[11], [12]. When the assumption of a scale-free topology is discarded, alternative clustering methods become more practical, as many graph clustering techniques relying on scale-freeness exploit the sparsity of edges between nodes[13]. Consequently, employing such methods without the scale-free assumption becomes computationally infeasible within a reasonable timeframe. One clustering method that does not rely on the scale-free assumption is spectral clustering (SC). Although SC has been studied for several decades, it continues to garner significant interest[14], [15]. SC has been applied in various fields, including biology, where it has been used to select optimal kernels for scRNA-seq data[16], and to classify structural domains in proteins[17]. The second limitation of WGCNA lies in the complexity of parameter tuning. Two essential parameters that must be manually determined, with the second parameter in particular requiring considerable effort to optimise. The third limitation is the disregard for regression coefficients. Even when a gene exhibits minimal variation in expression, a high correlation coefficient may still falsely imply co-expression.

To address these limitations, we propose a novel co-expression analysis method, SGCRNA (Spectral-clustering Generalised Correlation Regression Network Analysis).

## Materials and Methods

### datasets

Publicly available datasets were utilised for analysis (bulk RNA-Seq: GSE114007; Visium: GSE261545; metagenome: PRJNA762466).

### bulkRNA analysis

Fastp (v0.23.4) was employed to remove adapters and to filter raw reads shorter than 60 bases, as well as leading and trailing bases with a quality score below 30. Pseudoalignment of the filtered reads to the GRCh38 reference genome was performed using Kallisto (v0.48.0). GeTMM normalisation was performed using edgeR (v4.0.3) to account for inter-sample variation.

### Visium analysis

The R package Seurat was employed for spatial transcriptomic analysis. Specifically, raw data read via Read10X were processed using CreateSeuratObject (min.cells = 5, min.features = 5), followed by the estimation of doublet rates with DoubletFinder. Cells with mitochondrial gene content exceeding 10% were excluded as presumed dead cells. Subsequently, SCTransform, FindVariableFeatures (nfeatures = 3000), RunPCA, FindNeighbors (reduction = “pca”), and RunUMAP were performed sequentially.

### metagenome analysis

Metagenomic analysis was conducted using QIIME 2 (amplicon:2024.5), with taxonomic classification performed using the SILVA reference database (silva-138-99-nb-classifier.qza). Specifically, FASTQ files were imported using tools import, and PCR error correction was performed with dada2 denoise-paired (--p-trunc-len-f 297 --p-trunc-len-r 300). Taxonomic classification was subsequently conducted using feature-classifier classify-sklearn.

Low-read samples and Operational Taxonomic Units (OTUs) were excluded using feature-table filter-samples (--p-min-frequency 1500) and feature-table filter-features (--p-min-frequency 1), respectively. Taxa collapse was then applied to integrate data at the genus level, followed by normalisation using feature-table relative-frequency. Furthermore, prior to genus-level integration, OTUs corresponding to Faecalibacterium, Fusicatenibacter, or Eubacterium_coprostanoligenes_group were individually extracted, and gene function prediction was performed using PICRUSt2. The resulting data were visualised using KEGG Mapper, enabling the identification of pathways in which these genes are involved.

### WGCNA

The R package WGCNA was applied to both bulk RNA-Seq and 16S amplicon sequencing datasets. Specifically, the results of pickSoftThreshold were plotted, and the smallest power value exceeding the threshold of 0.9 was selected. The scale-free topology was verified by plotting the results obtained from adjacency. Subsequently, the Topological Overlap Matrix (TOM) was computed using TOMsimilarity, and hierarchical clustering was performed with hclust (method = “average”) using 1-TOM as the distance matrix. Module assignment was conducted with cutreeDynamic (DeepSplit = 4, MinModuleSize = 30), and module eigengenes were obtained using moduleEigengenes. Hierarchical clustering (method = “average”) was then performed on the module eigengenes, followed by module merging with mergeCloseModules (cutHeight = 0.3). Parameters not specifically mentioned were used at their default values.

For Visium data, the R package hdWGCNA was utilised. Specifically, SetupForWGCNA (gene_select = “fraction”, fraction = 0.05) was applied to the Seurat object. Subsequently, co-expression network analysis was conducted using SetDatExpr, TestSoftPowers, and ConstructNetwork.

## Results & Discussion

### Overview of SGCRNA workflows

SGCRNA is a novel method for the analysis of co-expression networks, grounded in correlation and linear relationships, and independent of specific network topologies (Fig. 1A). This method is applicable not only to transcriptomic data, but also to metagenomic, proteomic, and metabolomic variables, and accommodates a wide range of dataset types, including standard samples, time-course data, single-cell data, and spatially resolved datasets.

**Fig. 1.**
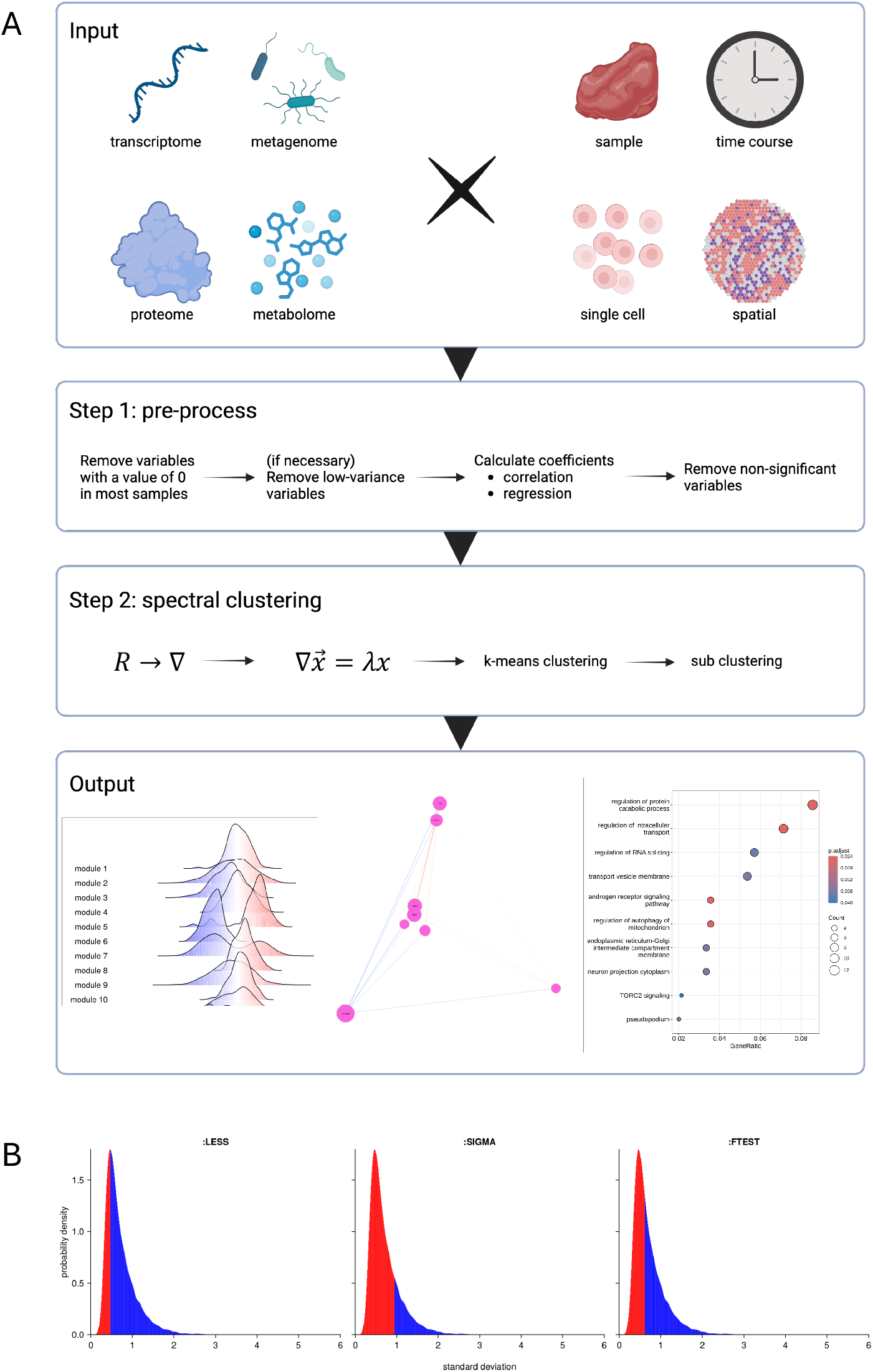
Overview of SGCRNA (A) Workflow of SGCRNA. (B) Optional pre-processing feature for removing low-variance variables and the corresponding exclusion ranges. From left to right: variables with standard deviations below the peak of the distribution (:LESS), variables with standard deviations within twice the peak standard deviation (:SIGMA), and variables with significantly lower standard deviations as determined by an F-test (:FTEST). Each panel depicts the distribution of standard deviations, with the x-axis representing standard deviation values and the y-axis indicating the number of variables exhibiting those values. The areas shaded in red represent the range of standard deviations corresponding to variables removed under each respective mode.

SGCRNA conducts analysis through a two-step process:

- Step 1: Data Pre-processing
  1. Variables with zero values in more than a specified proportion of the data (default: 50%) are removed.
  2. Low-variance variables may optionally be excluded (Fig. 1B).
    ⋄ Based on the variance distribution, three exclusion modes are available: below the peak (:LESS), within two standard deviations, assuming the peak corresponds to one sigma (:SIGMA), and within a range deemed significantly low variance by an F-test (:FTEST).
  3. Correlation coefficients and regression coefficients are then computed.
  4. Non-significant variables are excluded from further analysis.
- Step 2: Spectral Clustering
  1. When the correlation coefficient is represented as *cor* and the regression coefficient as *grad*, the edge weight is determined by the following equation:

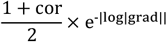
  2. Clustering is performed using a spectral clustering algorithm.
    ⋄ The weight matrix is subsequently transformed into a Laplacian matrix.
    ⋄ Eigenvectors are computed from the Laplacian matrix.
    ⋄ k-means clustering is then applied to the selected eigenvectors.

By calculating the correlations between modules derived from SGCRNA and traits, trait-associated modules can be identified. Moreover, by focusing on the scores assigned to nodes within each module (degree centrality adjusted by edge weights), hub variables exhibiting strong correlations with other variables can be identified. In the case of transcriptomic data, the node identifiers of each module can be retrieved for Gene Ontology (GO) analysis.

### SGCRNA outperforms existing methods in transcriptomic analysis

Osteoarthritis (OA) is known to become more prevalent with age and to occur more frequently in women than in men[18]. These tendencies were likewise observed in the dataset used in this study (Fig. 2A). Using this dataset, the most widely adopted gene co-expression network analysis method, WGCNA, was compared with SGCRNA. For WGCNA, parameters validating the scale-free topology assumption were applied (Supp. Fig. 1). SGCRNA partitioned the data into 76 modules, whereas WGCNA yielded only 14 (Fig. 2B). A notable characteristic of WGCNA clustering was the concentration of genes within specific modules. In contrast, SGCRNA yielded a more even distribution of gene counts across modules. Furthermore, module 73 of SGCRNA included genes that partially overlapped with both the magenta and grey modules of WGCNA, indicating distinct gene compositions between the two methods.

**Fig. 2.**
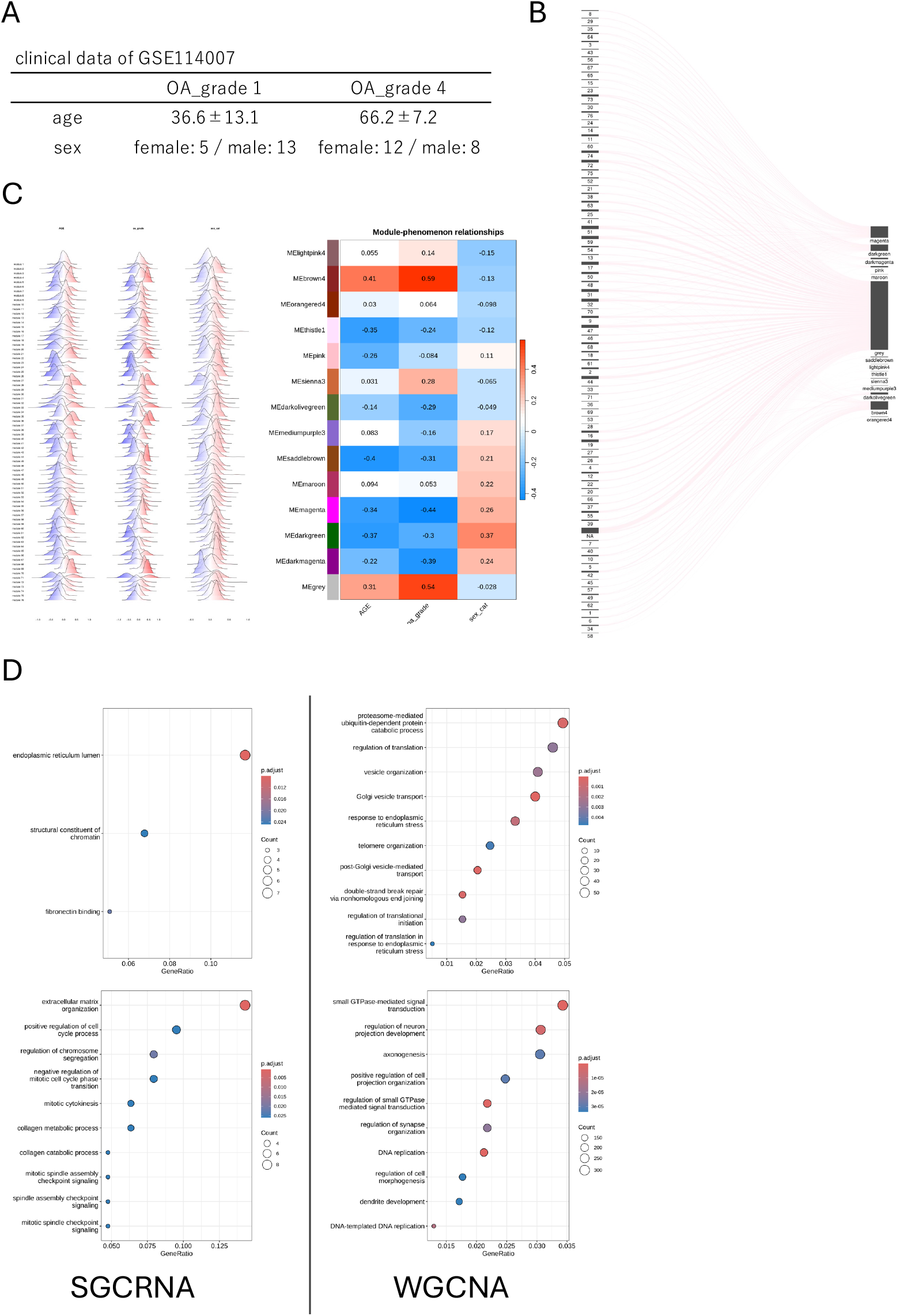
Transcriptomic Data Analysis Results by SGCRNA (A) Summary table of clinical data, including age, sex, and OA grade associated with the dataset. (B) S Sankey plot comparing clustering results between SGCRNA and WGCNA. (C) Correlations between modules and clinical data. Left: SGCRNA results—distributions of correlation coefficients between individual gene expression levels within each module and clinical data. Right: WGCNA results—heatmaps of correlation coefficients between module eigengenes and clinical data. (D) GO analysis of genes within modules associated with OA. Bubble plots for SGCRNA (top to bottom: module 45 and module 70) on the left; for WGCNA (top to bottom: module brown4 and module grey) on the right.

As previously noted, OA grade correlates positively with age and negatively with sex (coded as male = 0, female = 1). Therefore, correlations between module gene expression levels derived from both SGCRNA and WGCNA, and OA grade, age, and sex were examined (Fig. 2C). The results revealed that SGCRNA modules 45 and 70, and WGCNA modules brown4 and grey, were positively correlated with OA grade and age, and negatively correlated with sex. Subsequently, GO analysis was then performed on genes within these modules (Fig. 2D). In SGCRNA, module 45 comprised genes associated with the endoplasmic reticulum (ER), while module 70 included genes related to the extracellular matrix (ECM). Conversely, WGCNA’s brown4 module included genes linked to the Golgi apparatus, whereas the grey module encompassed a wide variety of ontologies. ER stress has been implicated in the pathological changes associated with OA[19], [20]. The ECM-related genes within SGCRNA module 70 included MMP11 and MMP13, as well as ADAMTS7 and ADAMTS14, which encode ECM-degrading enzymes implicated in OA progression[21]. Additionally, LAMB1 and LAMB3, components of the basement membrane known to be upregulated in OA[22], [23], along with FNDC1, a gene associated with the ECM and likewise upregulated in OA[24], [25], were identified. Although Golgi dysfunction has been suggested to contribute to OA pathogenesis[26], the broad range of ontologies encompassed by the grey module complicates interpretation.

Thus, SGCRNA offers the advantage of mitigating gene over-enrichment within specific modules and, by generating a greater number of finer clusters than WGCNA, facilitates more interpretable module analysis.

### SGCRNA also enables cell type-specific clustering in spatial transcriptomic data

This was validated using Visium data derived from kidney biopsies of patients with chronic kidney disease caused by diabetic and hypertensive nephropathy. Cell type classification within the tissue sections was referenced from the original study[27]. SGCRNA was compared with hdWGCNA, an adaptation of WGCNA suitable for high-dimensional data such as scRNA-Seq and Visium, using this dataset (Fig. 3). According to the original study, cell types for each spot were visually identified by a pathologist (Fig. 3A). SGCRNA partitioned the data into 114 modules, whereas hdWGCNA identified only two: grey and turquoise (Fig. 3B). Notably, the majority of genes were assigned to the grey module in hdWGCNA, approximately half of which had been excluded during SGCRNA pre-processing. When the expression distributions of genes within each module were plotted across the tissue sections, the grey module in hdWGCNA corresponded to tumour regions, while the turquoise module was associated with healthy proximal tubules (Fig. 3C). In contrast, while approximately 71% of SGCRNA-derived modules did not exhibit cell type-specific expression patterns, the remaining modules displayed specificity not only to tumour and healthy proximal tubules, but also to glomeruli, LH-CD, and artery regions (Fig. 3D, E). Although the original study identified additional cell types such as Inj-T, Cast-T, TLS, I-IFTA, and Capsule, SGCRNA-derived modules did not exhibit specificity for these cell types. This may be attributed to the gene filtering step in SGCRNA pre-processing, whereby variables expressed in fewer than half of the samples (in this case, spots) were excluded. Consequently, it is likely that genes expressed exclusively in cell types represented by a very small number of spots were removed during pre-processing. However, the spatially resolved enrichment patterns observed in SGCRNA-derived modules suggest that SGCRNA can still effectively capture cell type-specific transcriptional heterogeneity, which may be overlooked by conventional clustering approaches.

**Fig. 3.**
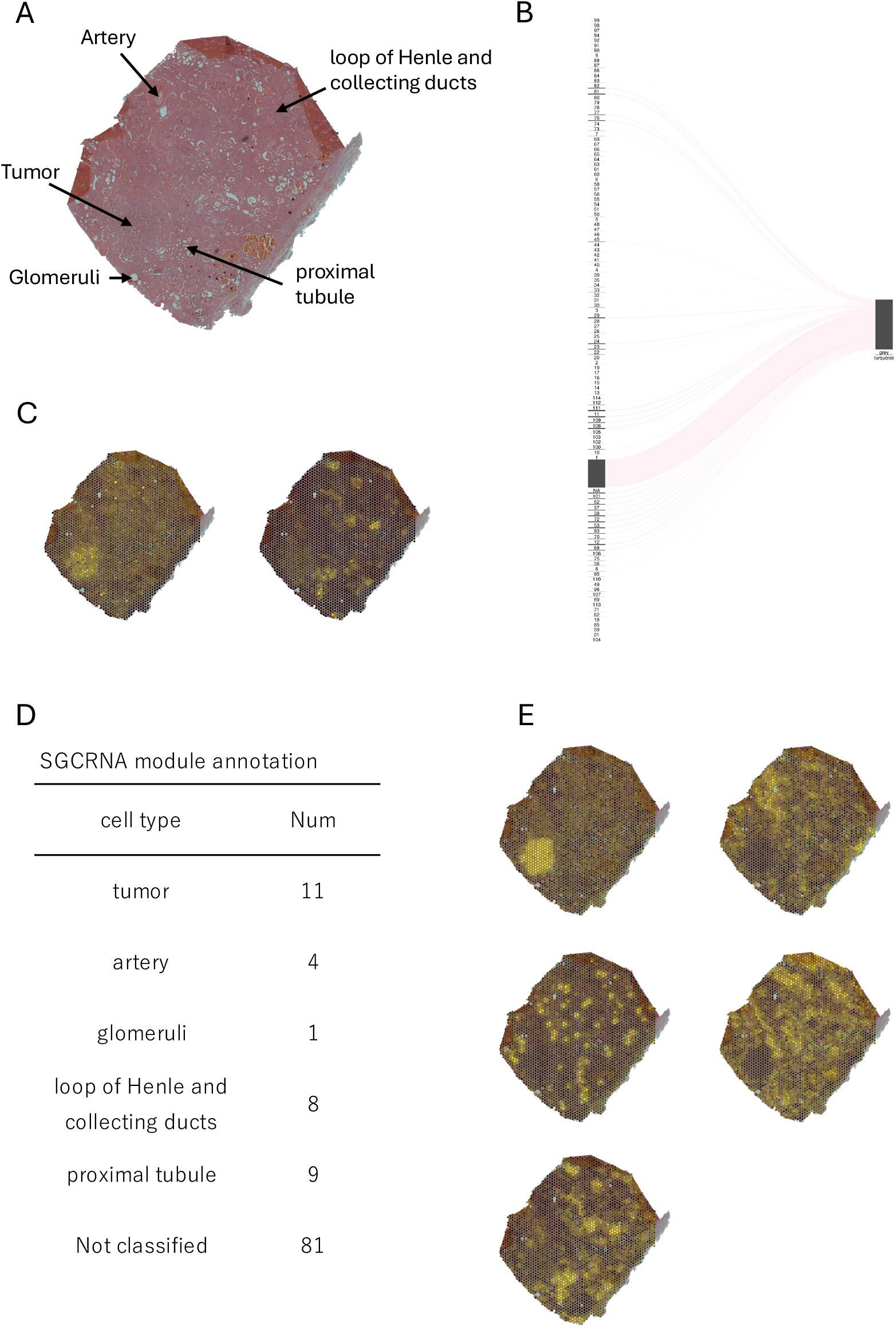
Spatial Transcriptomic Data Analysis Results by SGCRNA (A) Tissue section of the dataset and corresponding cell types. (B) Sankey plot comparing clustering results between SGCRNA and hdWGCNA. (C) Expression distribution of genes within each module identified by hdWGCNA. (D) Cell types in which gene expression within each SGCRNA-derived module was enriched, plotted onto tissue section images, along with the number of corresponding modules for each cell type. (E) Examples of gene expression distribution plots for SGCRNA-derived modules. Top row (left to right): tumour, artery; middle row (left to right): glomeruli, loop of Henle and collecting ducts; bottom row: proximal tubule.

### SGCRNA is also applicable to metagenomic datasets

This applicability was validated using 16S amplicon sequencing data derived from the human gut microbiota. Only OTUs identified at the genus level through taxonomic classification were included in the analysis; however, OTUs classified as “uncultured” or “Incertae Sedis” at the genus level were excluded. It should be noted that, unlike shotgun sequencing, 16S amplicon sequencing typically offers resolution at the family or genus level, rendering species-level identification challenging. A total of 208 genera were identified, and this dataset was analysed using WGCNA for comparative purposes. For WGCNA, no parameters yielding an adequate scale-free topology could be identified (Supp. Fig. 2). Following SGCRNA pre-processing, 15 genera were ultimately included in the network analysis (Fig. 4A). SGCRNA partitioned these into 12 modules, whereas WGCNA yielded only 8 (Fig. 4B). Nearly all of the 15 genera processed by SGCRNA were classified into the grey module by WGCNA, except for *Eubacterium coprostanoligenes group*, which was assigned to the red module. In contrast, SGCRNA clustered *Faecalibacterium* (Fa), *Fusicatenibacter* (Fu), and Eubacterium coprostanoligenes group (Eu) within a single module.

**Fig. 4.**
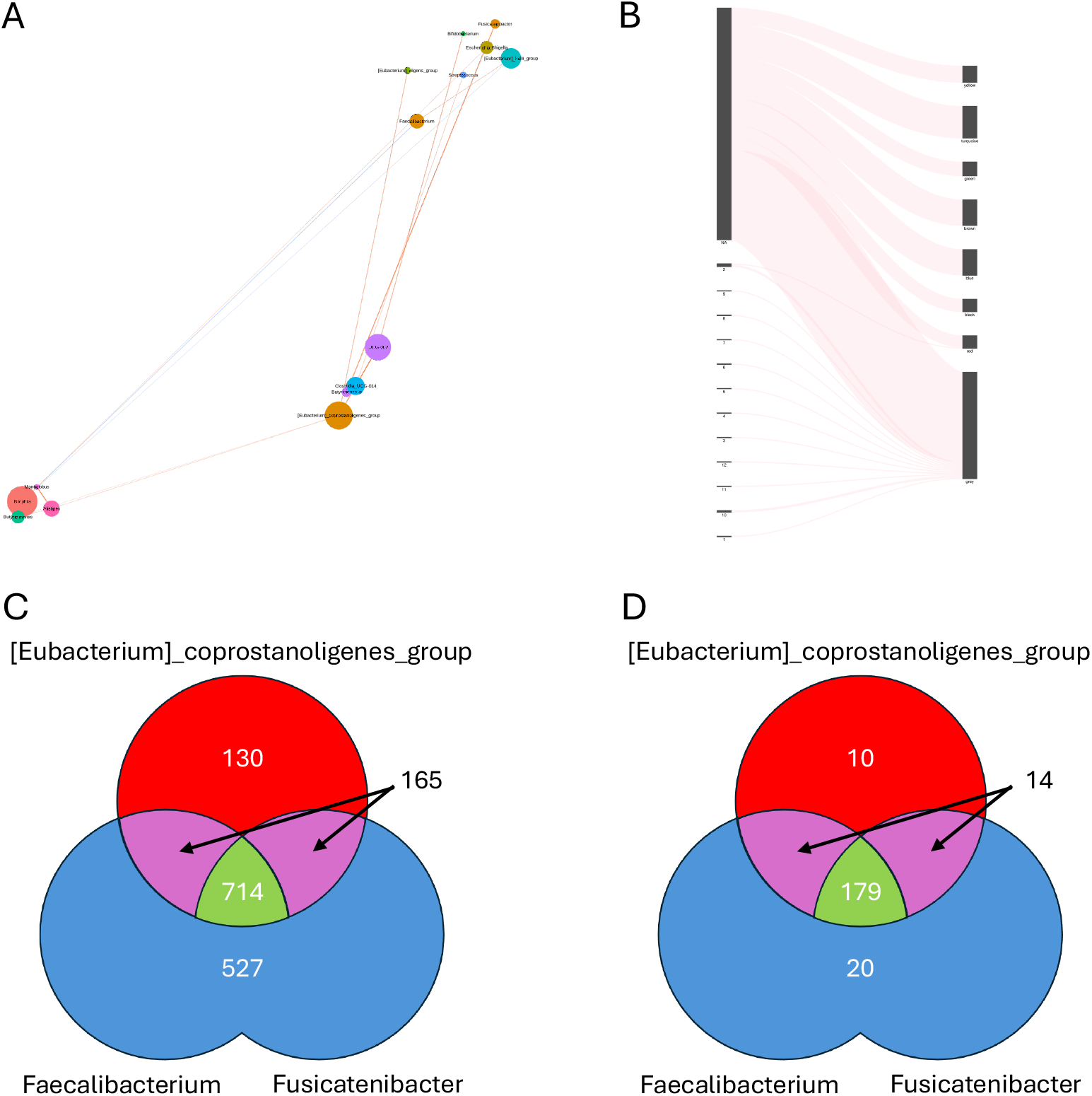
Metagenomic Data Analysis Results by SGCRNA (A) Co-expression network of gut microbiota. (B) Sankey plot comparing clustering results between SGCRNA and WGCNA. (C) Venn diagram showing the types of gene functions predicted by PICRUSt2.(D)Venn diagram showing the types of biological pathways associated with the gene functions predicted by PICRUSt2.

*Faecalibacterium* is a butyrate-producing bacterium regarded as playing a vital role in maintaining human health[28]. It has also been reported to exhibit significant inhibitory activity against α-amylase[29], thereby delaying carbohydrate digestion and absorption, which leads to reduced postprandial blood glucose levels. However, *Faecalibacterium* itself struggles to grow when utilising starch as a nutrient source[30].

*Fusicatenibacter* was named for its capacity to “devour sugars,” reflecting its possession of enzymes capable of polysaccharide degradation[31]. It has also been implicated in bile acid metabolism[32]. The Eubacterium coprostanoligenes group is known to degrade cholesterol, producing coprostanol as a metabolic by-product[33]. While bacteria are known to engage in cross-feeding—exchanging metabolic products as energy sources and nutrients across species or strains[34]—limited knowledge exists regarding potential symbiotic relationships among Fa, Fu, and Eu.

To investigate the functional relationships among these three genera, gene function prediction was conducted using PICRUSt2 (Fig. 4C). A total of 527 gene functions were identified within 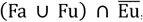, of which 125 were specific to Fa, 240 were specific to Fu, and 162 were shared between Fa and Fu. Additionally, 130 gene functions were specific to Eu, while 714 gene functions were common to all three genera. Moreover, 165 gene functions were identified in Eu ∩ (Fa Δ Fu). Based on these gene functions, KEGG Mapper was utilised to investigate the biological pathways associated with these genes (Fig. 4D). The results revealed 20 pathways corresponding to 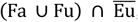,including 5 pathways specific to Fa, 9 specific to Fu, and 6 shared between Fa and Fu. Ten pathways were specific to Eu, while 179 pathways were common to all three genera. Additionally, 14 pathways were identified within Eu ∩ (Fa Δ Fu).

Gene function prediction using PICRUSt2 revealed that both Fu and Eu possess genes homologous to α-amylase among the three genera. With regard to glycolysis, the function of extracellular glucose uptake was predicted for Fa and Fu, but not for Eu. Specifically, among the gene functions associated with Enzyme Commission number (EC) 2.7.1.199—K02777, K02779, K02791, and K20118—K02779 was not predicted by PICRUSt2. K02777 and K02791 were predicted in Fa, whereas K20118 was predicted in both Fa and Fu. Although Eu is known to utilise glucose as a nutrient, this is likely mediated by the ABC transporter gene K10112, which was predicted in all three genera. Therefore, glucose uptake in Eu is presumed to be mediated by K10112. The butyrate production pathway was predicted in Fa and Fu, but was largely absent in Eu. Extracellular glucose is also utilised in the synthesis of GlcNAc-1-phosphate, a precursor in the peptidoglycan biosynthesis pathway essential for bacterial cell wall formation. I In the fatty acid degradation pathway, functional divergence was observed. For the conversion of fatty acids to aldehydes (EC 1.2.1.3), among the related genes K00128, K14085, and K00149, only K00128 was predicted, and it was specific to Fu. Regarding the conversion of aldehydes to alcohols (EC 1.1.1.1), among K13951, K13980, K13952, K13953, K13954, K00001, K00121, K04072, and K18857, only K13954 (in Fu), and K00001 and K04072 (in Eu) were predicted. Alcohol can subsequently be further metabolised via glycolytic pathways.

Based on these findings, the following hypothesis is proposed (Supp. Fig. 3). Fa utilises glucose, which is derived from host amylase activity and the enzymatic breakdown by Fu and Eub, as a nutrient source to produce butyrate and peptidoglycan. In the presence of Fu and Eu, bile acids are metabolised by Fu into aldehydes, which, due to the anaerobic environment, diffuse and are subsequently converted into alcohols by Eu. The alcohols, in turn, diffuse and are taken up by Fa, where they are metabolised through butanonate metabolism to produce butyrate. Thus, it is hypothesised that butyrate production in this microbial consortium occurs not only via the conventional glucose-derived pathway, but also through a bile acid-derived pathway, thereby contributing to the maintenance of gut homeostasis. Although this hypothesis is supported by predicted gene functions, it requires experimental validation. Further *in vitro* or *in vivo* investigations are necessary to substantiate the presence of such cross-feeding interactions and their contribution to butyrate synthesis.

### Limitations

As SGCRNA employs spectral clustering, the computation of eigenvectors is required. The computational complexity of calculating the eigenvectors of a square matrix of order *n* is generally *O*(*n*^3^). However, SGCRNA utilises an approximate method based on the Krylov subspace, reducing the complexity to *O*(*k* * *n*^2^), where k is the number of eigenvectors to be computed. Consequently, a notable limitation of SGCRNA is that computational time increases significantly with the number of variables. Indeed, when compared to WGCNA, SGCRNA requires substantially more execution time (Supp. Fig. 4). Execution time comparisons were conducted using dataset GSE114007 on an Intel Xeon Gold 6230 processor, with each method executed 10 times. For WGCNA, no parameter tuning was performed; instead, arbitrary values were employed (power = 20, cutHeight = 0.9). Moreover, since WGCNA typically requires visual inspection of plots for parameter selection, the time required to generate these plots was included in the measurements. It is important to note that WGCNA is implemented as an R package, whereas SGCRNA is developed in Julia, and thus direct comparisons of execution speed should be interpreted with caution. However, considering that WGCNA leverages C-compiled code for faster execution relative to native R code, and that Julia, when compiled, generally exhibits execution times approximately 2 to 5 times slower than C++ (whereas R may be hundreds of times slower)[35], these results may be considered a general indication of relative performance.

## Data availability

The authors confirm that the data supporting the findings of this study are available within the article.

## Acknowledgments

Grants-in-Aid for Scientific Research from the Japan Society for the Promotion of Science (23K14384 to T. Osone; 23K08677 to T. Takao; 23K21368 to T. Takarada), AMED (JP24bm1123059 to T.Takao), and JST FOREST Program (JPMJFR225H to T. Takarada). These funders had no role in the study design, data collection and analysis, decision to publish, or preparation of the manuscript.

The computation was carried out using the General Projects on supercomputer “Flow” at Information Technology Center, Nagoya University.

## Figure legends

Sup. Fig. 1

WGCNA Parameter Tuning Results for Transcriptomic Data. Top left to bottom right: determination of the soft threshold, evaluation of scale-free topology at the selected soft threshold, clustering results by Dynamic Tree Cut.

Sup. Fig. 2

WGCNA Parameter Tuning Results for Metagenomic Data. Top left to bottom right: determination of the soft threshold, evaluation of scale-free topology at the selected soft threshold, clustering results by Dynamic Tree Cut.

Sup. Fig. 3

Cross-feeding Hypothesis Diagram for Fa, Fu, and Eu. Coloured boxes represent EC numbers or biological pathways. Red: Eu-specific gene functions; Blue: gene functions in 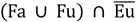; Purple: gene functions in Eu ∩ (Fa Δ Fu); Green: gene functions common to all three genera.

Sup. Fig. 4

Execution Time Comparison between WGCNA and SGCRNA. Analysis time measured using GSE114007 with varying numbers of variables. X-axis: number of variables (genes); Y-axis: execution time. Red triangles represent SGCRNA results; blue crosses represent WGCNA results.

## References

[1] B. Zhang and S. Horvath, ‘A General Framework for Weighted Gene Co-Expression Network Analysis’, Stat Appl Genet Mol Biol, vol. 4, no. 1, Jan. 2005, doi: 10.2202/1544-6115.1128.

[2] S. Zheng and W.-X. Wang, ‘Differential effects of foodborne and waterborne micro(nano)plastics exposure on fish liver metabolism and gut microbiota community’, J Hazard Mater, vol. 488, p. 137471, May 2025, doi: 10.1016/j.jhazmat.2025.137471.

[3] Y. Zhou et al., ‘Prediction of influenza virus infection based on deep learning and peripheral blood proteomics: A diagnostic study’, J Adv Res, Mar. 2025, doi: 10.1016/j.jare.2025.03.051.

[4] M. Kouhsar et al., ‘A brain DNA co-methylation network analysis of psychosis in Alzheimer’s disease’, Alzheimer’s & Dementia, vol. 21, no. 2, Feb. 2025, doi: 10.1002/alz.14501.

[5] N. Rezaie, F. Reese, and A. Mortazavi, ‘PyWGCNA: a Python package for weighted gene co-expression network analysis’, Bioinformatics, vol. 39, no. 7, Jul. 2023, doi: 10.1093/bioinformatics/btad415.

[6] S. García-Ruiz et al., ‘CoExp: A Web Tool for the Exploitation of Co-expression Networks’, Front Genet, vol. 12, Feb. 2021, doi: 10.3389/fgene.2021.630187.

[7] S. Morabito, F. Reese, N. Rahimzadeh, E. Miyoshi, and V. Swarup, ‘hdWGCNA identifies co-expression networks in high-dimensional transcriptomics data’, Cell Reports Methods, vol. 3, no. 6, p. 100498, Jun. 2023, doi: 10.1016/j.crmeth.2023.100498.

[8] Y. Liu, ‘CWGCNA : an R package to perform causal inference from the WGCNA framework’, NAR Genom Bioinform, vol. 6, no. 2, Apr. 2024, doi: 10.1093/nargab/lqae042.

[9] Y. Liu et al., ‘TPSC: a module detection method based on topology potential and spectral clustering in weighted networks and its application in gene co-expression module discovery’, BMC Bioinformatics, vol. 22, no. S4, p. 111, Oct. 2021, doi: 10.1186/s12859-021-03964-5.

[10] P. Langfelder and S. Horvath, ‘WGCNA: an R package for weighted correlation network analysis’, BMC Bioinformatics, vol. 9, no. 1, p. 559, Dec. 2008, doi: 10.1186/1471-2105-9-559.

[11] H. B. Smith, H. Kim, and S. I. Walker, ‘Scarcity of scale-free topology is universal across biochemical networks’, Sci Rep, vol. 11, no. 1, p. 6542, Mar. 2021, doi: 10.1038/s41598-021-85903-1.

[12] A. D. Broido and A. Clauset, ‘Scale-free networks are rare’, Nat Commun, vol. 10, no. 1, p. 1017, Mar. 2019, doi: 10.1038/s41467-019-08746-5.

[13] V. D. Blondel, J.-L. Guillaume, R. Lambiotte, and E. Lefebvre, ‘Fast unfolding of communities in large networks’, Journal of Statistical Mechanics: Theory and Experiment, vol. 2008, no. 10, p. P10008, Oct. 2008, doi: 10.1088/1742-5468/2008/10/P10008.

[14] F. Nie, J. Xue, R. Wang, L. Zhang, and X. Li, ‘Fast Clustering by Directly Solving Bipartite Graph Clustering Problem’, IEEE Trans Neural Netw Learn Syst, vol. 35, no. 7, pp. 9174–9185, Jul. 2024, doi: 10.1109/TNNLS.2022.3219131.

[15] R. Wang, H. Chen, Y. Lu, Q. Zhang, F. Nie, and X. Li, ‘Discrete and Balanced Spectral Clustering With Scalability’, IEEE Trans Pattern Anal Mach Intell, vol. 45, no. 12, pp. 14321–14336, Dec. 2023, doi: 10.1109/TPAMI.2023.3311828.

[16] R. Qi, J. Wu, F. Guo, L. Xu, and Q. Zou, ‘A spectral clustering with self-weighted multiple kernel learning method for single-cell RNA-seq data’, Brief Bioinform, vol. 22, no. 4, Jul. 2021, doi: 10.1093/bib/bbaa216.

[17] D. Granata, L. Ponzoni, C. Micheletti, and V. Carnevale, ‘Patterns of coevolving amino acids unveil structural and dynamical domains’, Proceedings of the National Academy of Sciences, vol. 114, no. 50, Dec. 2017, doi: 10.1073/pnas.1712021114.

[18] S. Glyn-Jones et al., ‘Osteoarthritis’, The Lancet, vol. 386, no. 9991, pp. 376–387, Jul. 2015, doi: 10.1016/S0140-6736(14)60802-3.

[19] Z. Wen et al., ‘Endoplasmic Reticulum Stress in Osteoarthritis: A Novel Perspective on the Pathogenesis and Treatment’, Aging Dis, p. 0, 2022, doi: 10.14336/AD.2022.0725.

[20] D. de Seny et al., ‘Proteins involved in the endoplasmic reticulum stress are modulated in synovitis of osteoarthritis, chronic pyrophosphate arthropathy and rheumatoid arthritis, and correlate with the histological inflammatory score’, Sci Rep, vol. 10, no. 1, p. 14159, Sep. 2020, doi: 10.1038/s41598-020-70803-7.

[21] A. Mukherjee and B. Das, ‘The role of inflammatory mediators and matrix metalloproteinases (MMPs) in the progression of osteoarthritis’, Biomaterials and Biosystems, vol. 13, p. 100090, Mar. 2024, doi: 10.1016/j.bbiosy.2024.100090.

[22] S. L. Dunn, J. Soul, S. Anand, J.-M. Schwartz, R. P. Boot-Handford, and T. E. Hardingham, ‘Gene expression changes in damaged osteoarthritic cartilage identify a signature of non-chondrogenic and mechanical responses’, Osteoarthritis Cartilage, vol. 24, no. 8, pp. 1431–1440, Aug. 2016, doi: 10.1016/j.joca.2016.03.007.

[23] R. Piva et al., ‘Slug transcription factor and nuclear Lamin B1 are upregulated in osteoarthritic chondrocytes’, Osteoarthritis Cartilage, vol. 23, no. 7, pp. 1226–1230, Jul. 2015, doi: 10.1016/j.joca.2015.03.015.

[24] P. Yi, X. Xu, J. Yao, and B. Qiu, ‘Effect of DNA methylation on gene transcription is associated with the distribution of methylation sites across the genome in osteoarthritis’, Exp Ther Med, vol. 22, no. 1, p. 719, May 2021, doi: 10.3892/etm.2021.10151.

[25] R. Bouchareb et al., ‘Proteomic Architecture of Valvular Extracellular Matrix’, JACC Basic Transl Sci, vol. 6, no. 1, pp. 25–39, Jan. 2021, doi: 10.1016/j.jacbts.2020.11.008.

[26] G. L. Iacobescu, A.-D. Corlatescu, M. Popa, L. Iacobescu, C. Cirstoiu, and C. Orban, ‘Exploring the Implications of Golgi Apparatus Dysfunction in Bone Diseases’, Cureus, Mar. 2024, doi: 10.7759/cureus.56982.

[27] P. Isnard, D. Li, Q. Xuanyuan, H. Wu, and B. D. Humphreys, ‘Histopathologic Analysis of Human Kidney Spatial Transcriptomics Data’, Am J Pathol, vol. 195, no. 1, pp. 69–88, Jan. 2025, doi: 10.1016/j.ajpath.2024.06.011.

[28] M. Lopez-Siles, S. H. Duncan, L. J. Garcia-Gil, and M. Martinez-Medina, ‘Faecalibacterium prausnitzii : from microbiology to diagnostics and prognostics’, ISME J, vol. 11, no. 4, pp. 841–852, Apr. 2017, doi: 10.1038/ismej.2016.176.

[29] X. Sun, Z. Zhang, and J. Hu, ‘Isolation, probiotic characterization and whole-genome sequencing of gut Faecalibacterium prausnitzii’, Human Nutrition & Metabolism, vol. 40, p. 200315, Jun. 2025, doi: 10.1016/j.hnm.2025.200315.

[30] M. Lopez-Siles, T. M. Khan, S. H. Duncan, H. J. M. Harmsen, L. J. Garcia-Gil, and H. J. Flint, ‘Cultured Representatives of Two Major Phylogroups of Human Colonic Faecalibacterium prausnitzii Can Utilize Pectin, Uronic Acids, and Host-Derived Substrates for Growth’, Appl Environ Microbiol, vol. 78, no. 2, pp. 420–428, Jan. 2012, doi: 10.1128/AEM.06858-11.

[31] T. Takada, T. Kurakawa, H. Tsuji, and K. Nomoto, ‘Fusicatenibacter saccharivorans gen. nov., sp. nov., isolated from human faeces’, Int J Syst Evol Microbiol, vol. 63, no. Pt_10, pp. 3691–3696, Oct. 2013, doi: 10.1099/ijs.0.045823-0.

[32] P. Gao et al., ‘Gut microbial metabolism of bile acids modifies the effect of Mediterranean diet interventions on cardiometabolic risk in a randomized controlled trial’, Gut Microbes, vol. 16, no. 1, Dec. 2024, doi: 10.1080/19490976.2024.2426610.

[33] A. Mukherjee, C. Lordan, R. P. Ross, and P. D. Cotter, ‘Gut microbes from the phylogenetically diverse genus Eubacterium and their various contributions to gut health’, Gut Microbes, vol. 12, no. 1, p. 1802866, Nov. 2020, doi: 10.1080/19490976.2020.1802866.

[34] E. J. Culp and A. L. Goodman, ‘Cross-feeding in the gut microbiome: Ecology and mechanisms’, Cell Host Microbe, vol. 31, no. 4, pp. 485–499, Apr. 2023, doi: 10.1016/j.chom.2023.03.016.

[35] J. Bezanson, S. Karpinski, V. B. Shah, and A. Edelman, ‘Julia: A Fast Dynamic Language for Technical Computing’, Sep. 2012.

