## Supplementary figures and images for "SGCRNA: Spectral Clustering-Guided Co-Expression Network Analysis Without Scale-Free Constraints for Multi-Omic Data"

### Sup. Fig. 1; Sup. Fig. 2; Sup. Fig. 3; Sup. Fig. 4

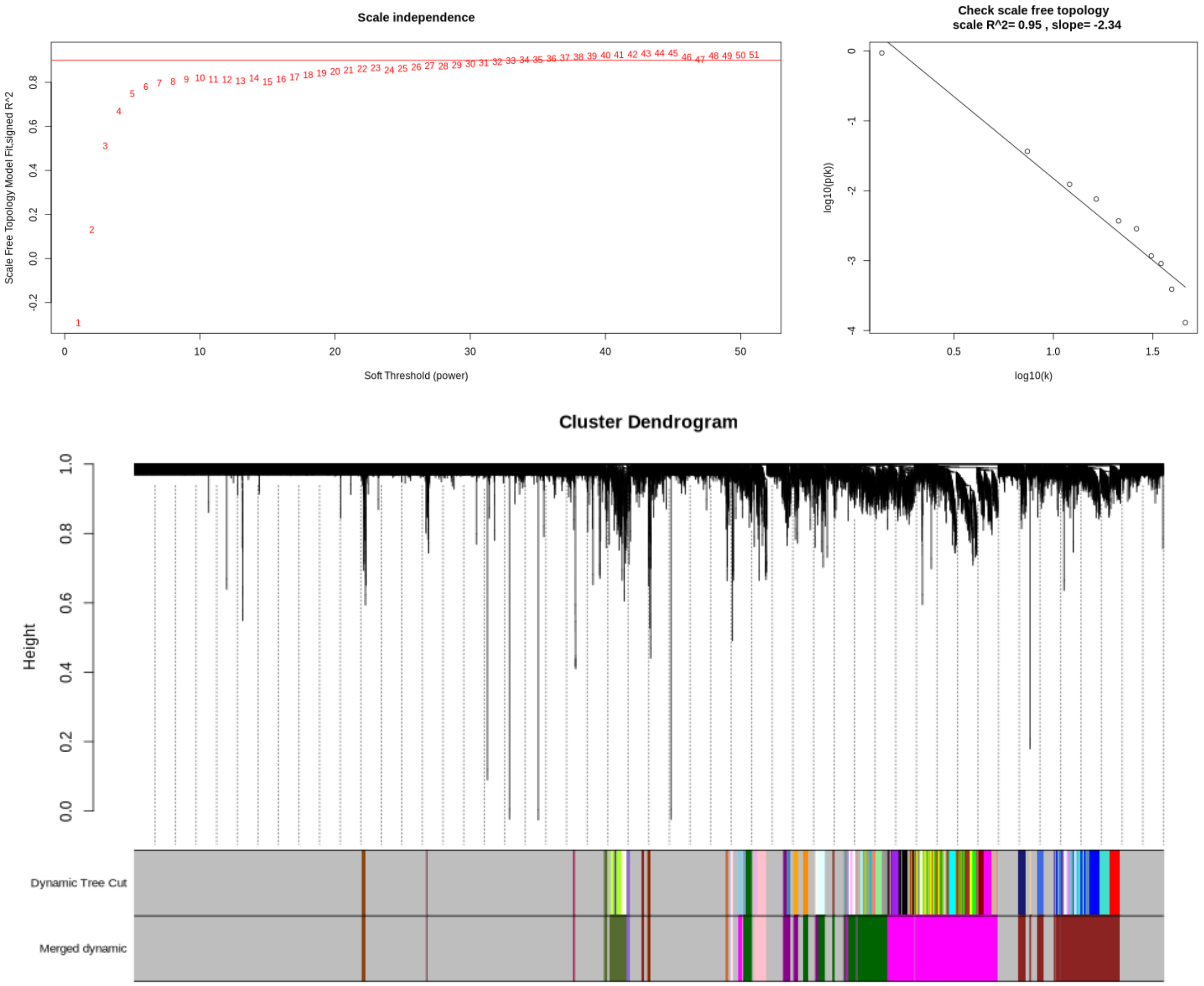

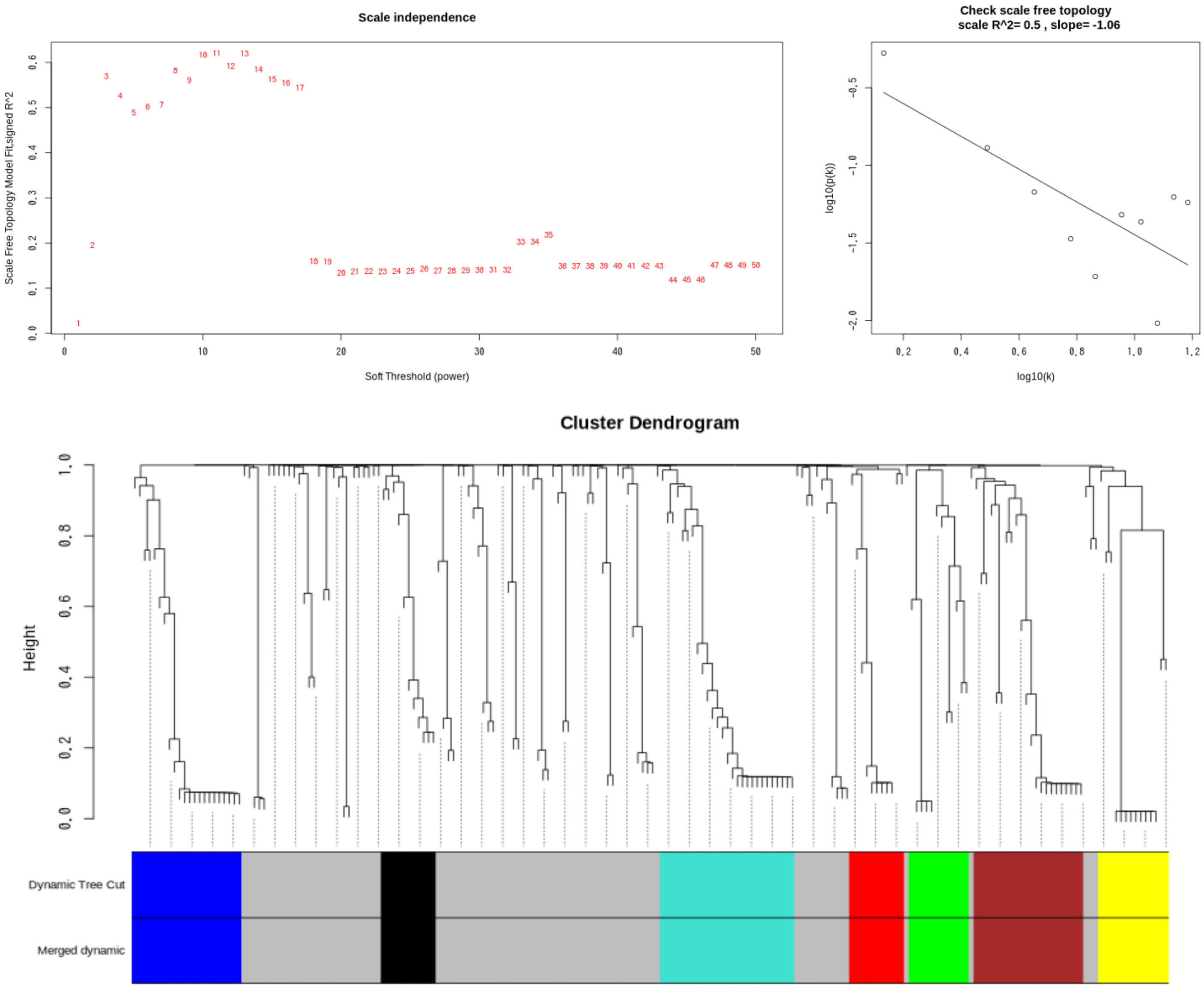


Sup. Fig. 2

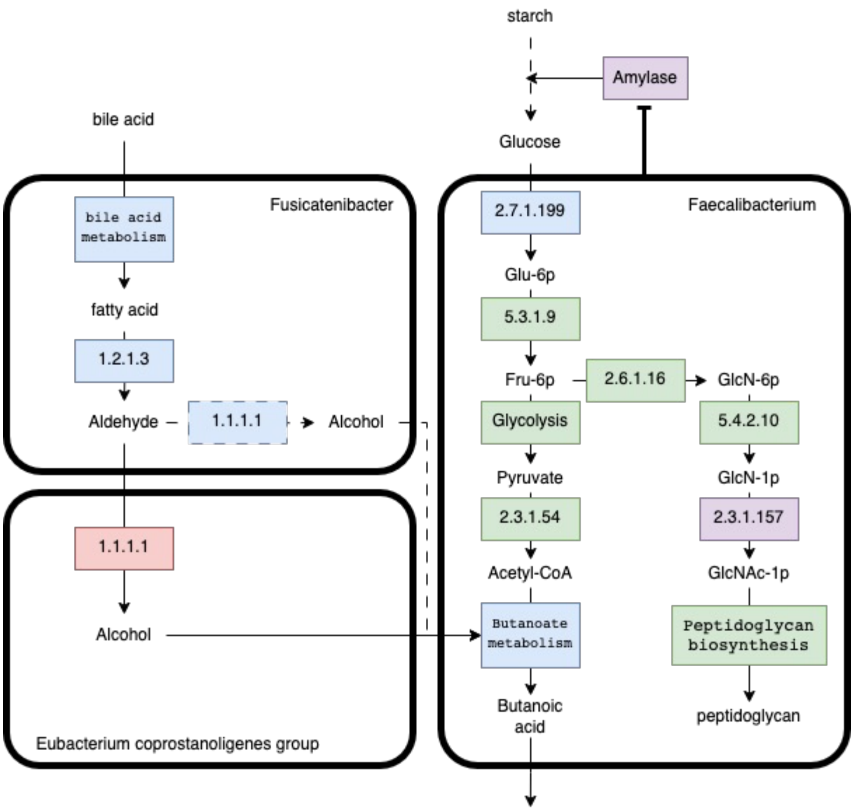


Sup. Fig. 3

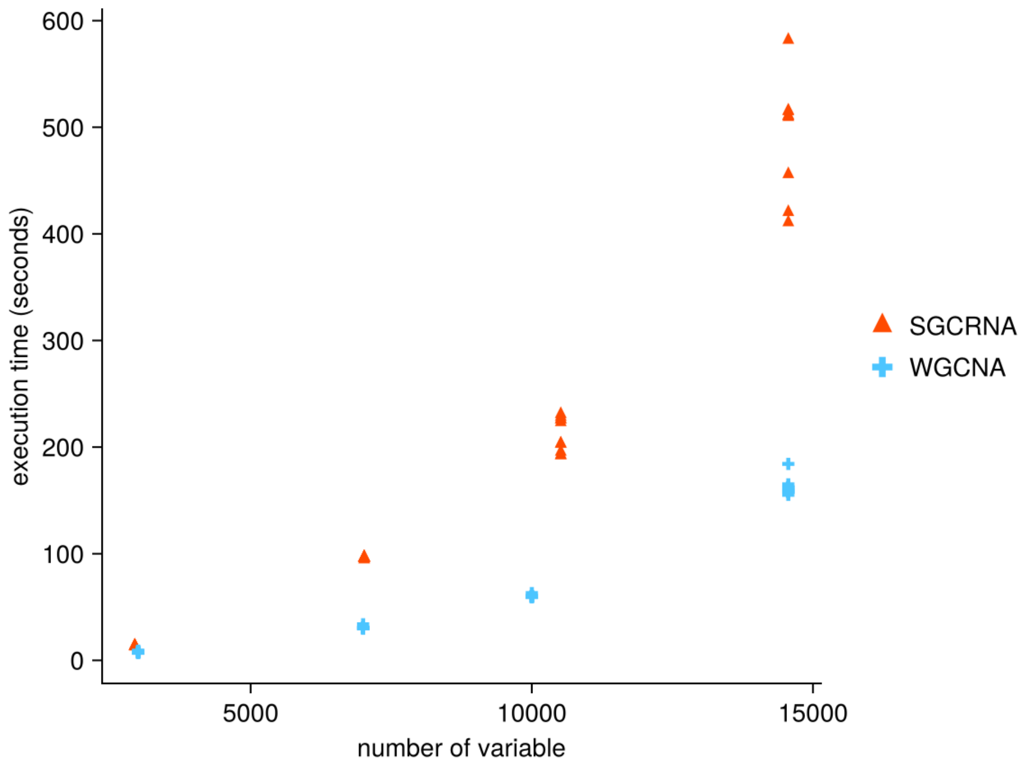
